## Supplementary material for "*Mycobacterium tuberculosis* H_2_S functions as a sink to modulate central metabolism, bioenergetics, and drug susceptibility": Mtb-H2S Supplementary Materials

**Supplementary Figures and Tables**

**Supplementary Figure 1.** *Mtb* H_2_S production when exposed to exogenous Cys.

**Supplementary Figure 2.** Role of *Mtb* Rv1077 (CBS) in H_2_S production.

**Supplementary Figure 3.** Multiple sequence alignment of the CDS/CBS protein family.

**Supplementary Figure 4.** Complete Rv3684/Cds1 amino acid sequence.

**Supplementary Figure 5.** In-gel BC assay of purified Cds1 and confirmation of the cds1 knockout mutant in *Mtb* strains.

**Supplementary Figure 6.** Survival of *cds1*-deficient *Mtb* in macrophages.

**Supplementary Figure 7.** *Mtb* Δ*cds1* growth in the presence of fatty acids or precursors as a single carbon source.

**Supplementary Figure 8.** *Mtb* H_2_S production when cultured in the presence of fatty acids or precursors as a single carbon source.

**Supplementary Figure 9.** Role of Cbs (Rv1077) in *Mtb* respiration.

**Supplementary Figure 10.** AOAA inhibits Cys-mediated increases in *Mtb* respiration.

**Supplementary Figure 11.** Exogenous H_2_S reverses the respiratory defect in *Mtb ∆cds1* cells.

**Supplementary Figure 12.** SDS-PAGE of purified *O*-acetylserine sulfhydrylase (OASS).

**Supplementary Figure 13.** Cds1 regulates amino acid metabolism in *Mtb*.

**Supplementary Figure 14.** Gating strategy for detection of DHE-positive *Mtb* cells for measuring ROI.

**Supplementary Figure 15.** Mycothiol and ergothioneine levels in *Mtb* after exposure to CHP.

**Supplementary Table 1:** *Mtb* H37Rv enzymes putatively involved in sulfur-containing amino acid biosynthesis, H_2_S production or sulfur metabolism.

**Supplementary Table 2*:*** Bacterial strains used in this study.

**Supplementary Table 3:** Plasmids used in this study.

**Supplementary Table 4:** Oligonucleotides used in this study.

**REFERENCES**

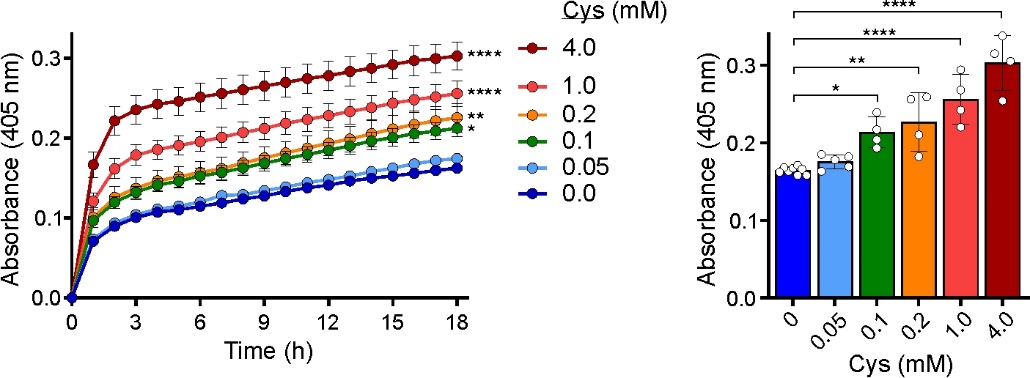

**Supplementary Figure 1. *Mtb* H_2_S production when exposed to exogenous Cys.** The BC assay was used to measure H_2_S production in *Mtb* when exposed to increasing concentrations of exogenous Cys (0-4 mM). The kinetics of H_2_S production is shown in the left panel, whereas endpoint levels after 18 hours are shown in the right panel. Note that *Mtb* does not require addition of exogenous Cys to produce H_2_S (see 0 mM Cys) and Fig. 1. Data shown represents the mean ± SEM for 4 – 5 biological replicates. Statistical analysis was performed using GraphPad Prism 7.02. Two-way ANOVA with Dunnett’s multiple comparisons test was used to determine statistical significance compared to 0.0 mM Cys. **P <* 0.05, ***P* < 0.01, *****P <* 0.0001.

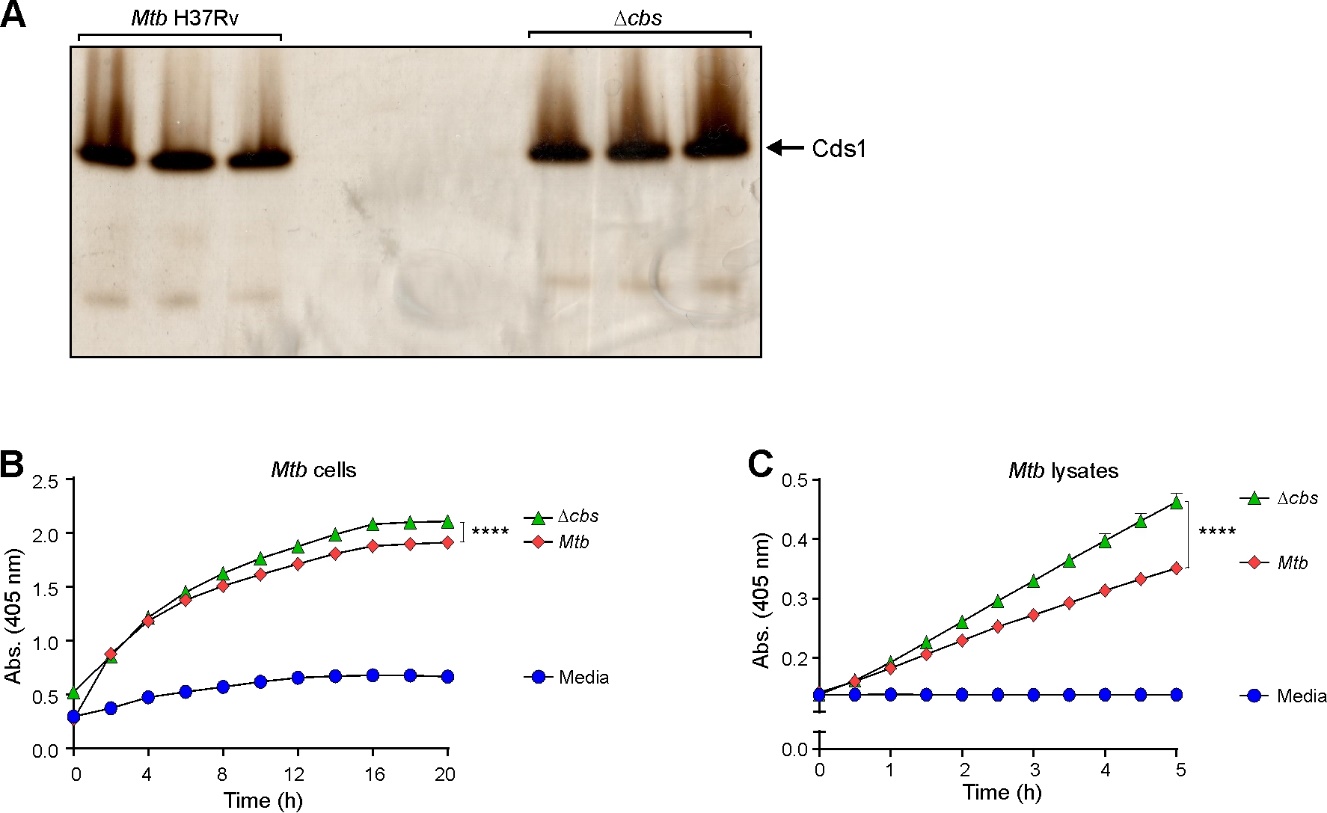

**Supplementary Figure 2. Role of *Mtb* Rv1077 (CBS) in H_2_S production. (A)** Lysates of *Mtb* H37Rv and the *Mtb* *rv1077*/*cbs* deletion strain (Δ*cbs*) were separated on a native polyacrylamide gel and assayed for H_2_S production using the in-gel BC assay. Arrow indicates the major H_2_S producing enzyme, (*n* = 3). Time course of H_2_S production in **(B)** intact *Mtb* H37Rv and Δ*cbs* cells, (*n* = 8) and **(C)** cell lysates in the presence of 20 mM Cys using the BC assay (*n* = 8). Representative experiments are shown. Each experiment was repeated independently at least twice. Data represent the mean ± SD. Statistical analysis was performed using GraphPad Prism 7.02. One-way ANOVA with Dunnett’s multiple comparisons test was used to determine statistical significance. *****P* < 0.0001.

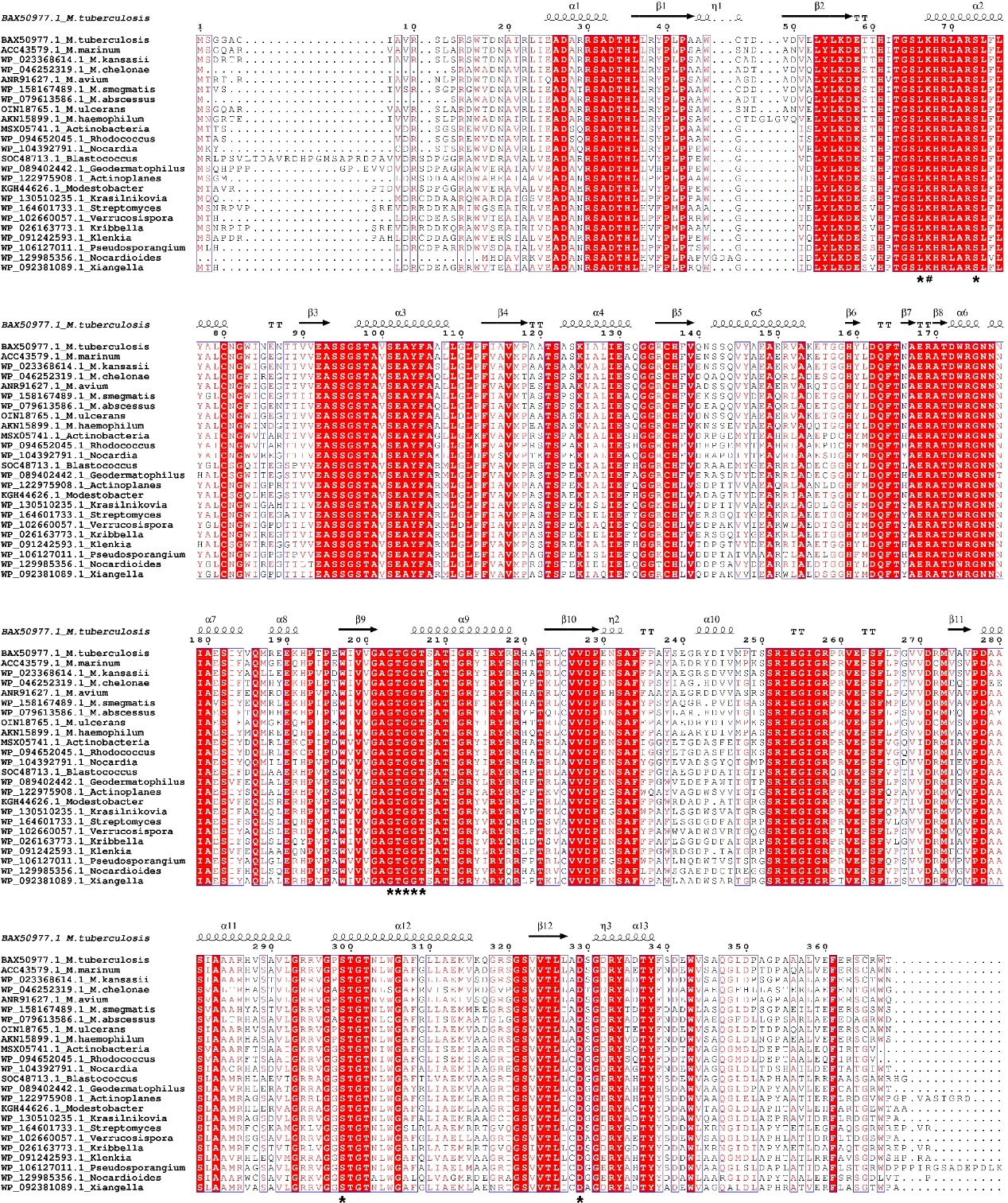
**Supplementary Figure 3.** **Multiple sequence alignment of the CDS/CBS protein family.** Multiple sequence alignment of Cds1 BLASTp. Structural alignment performed using T-Coffee online software^1^ and rendered with predicted secondary structures using the ESPript. 3.0 server^2^. Identical amino acids are shaded in red. Asterisks (*) indicate residues of the predicted, conserved PLP binding domain (L^66^, S^73^, G^203^, T^204^, G^205^, G^206^, T^207^, S^209^, D^329^). # indicates the predicted catalytic residue (K^67^) which is conserved across orthologues with an identity > 65%**.** Sequences were obtained from a BLASTp search of the NCBI database (https://blast.ncbi.nlm.nih.gov/Blast.cgi) using the primary sequence of Cds1 as identified by LC-MS/MS (Fig. 2c). CDS; cysteine desulfhydrase, CBS; cystathionine β-synthase.

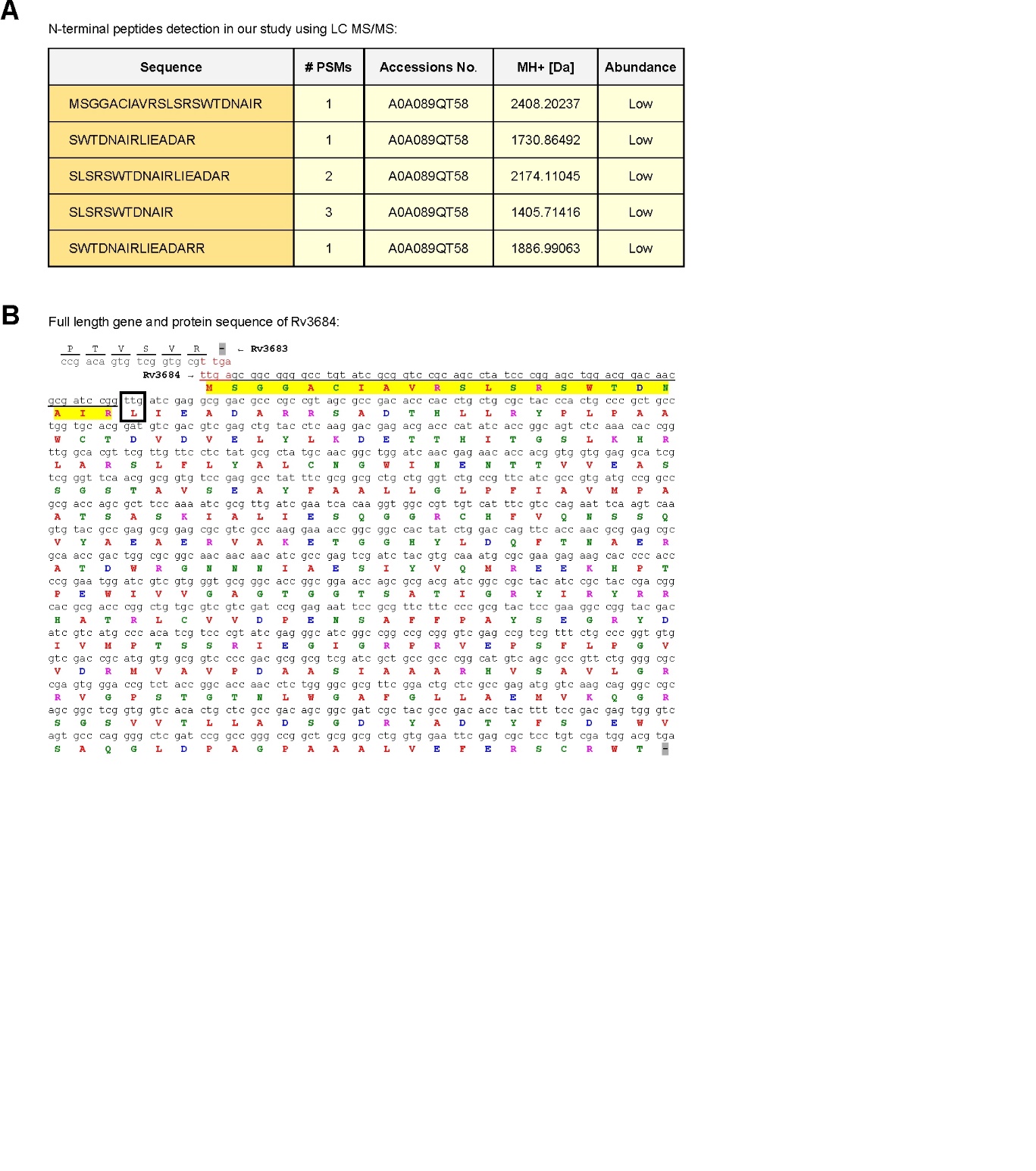

**Supplementary Figure 4. Complete Rv3684/Cds1 amino acid sequence. (A)** A list of N-terminal peptide fragments of Cds1 detected by LC-MS/MS corresponding to the correct amino acid sequence of Cds1. (**B**) The correct *rv3684/cde1* ORF includes an additional 66 nucleotides encoding 22 additional N-terminal amino acids (highlighted in yellow) in contrast to the annotation in Mycobrowser.epfl.ch. The start codon of *cds1* overlaps the stop codon of *rv3683* and both ORFs are in different coding frames. The predicted start codon (ttg) of *cds1* in Mycobrowser.epfl.ch is indicated in the box. Color code of amino acid residues: Hydrophobic - AFILMVW (red); Polar - CGHNPQSTY (green); Basic charged - K and R (pink); Acidic charged - D and E (blue).

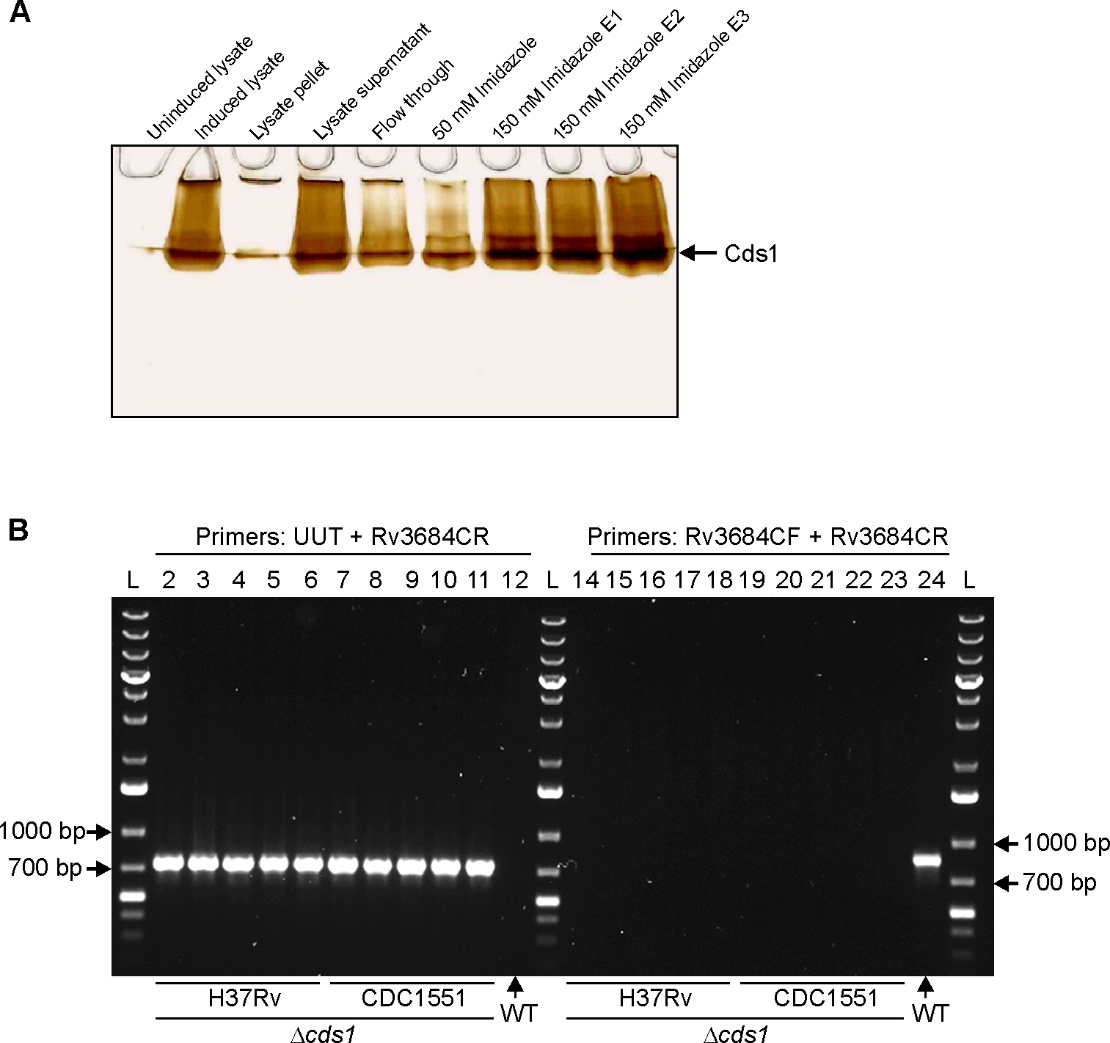

**Supplementary Figure 5. In-gel BC assay of purified Cds1 and confirmation of the cds1 knockout mutant in *Mtb* strains. (A)** Eluted fractions of recombinant Cds1 were resolved on a native polyacrylamide gel and assayed for H_2_S production using the in-gel BC assay. **(B)** PCR confirmation of *Mtb* H37Rv (*n* =5) and CDC1551 *cds1* knockout mutants (*n* = 5). Bands in lanes 2 to 6 (*Mtb* H37Rv) and lanes 7 to 11 (*Mtb* CDC1551) correspond to PCR amplicons (727 bp) generated using primers UUT and Rv3684CR. As the annealing site for primer UUT is only present in the disrupted *cds1*, no amplicon was generated for WT *Mtb* H37Rv (lane 12). The band in lane 24 corresponds to the PCR amplicon (819 bp) generated in WT *Mtb* H37Rv using primers Rv3684CF and Rv3684CR. The annealing site for primer Rv3684CF is present in *cds1,* but absent in the *cds1* mutant and therefore no PCR amplicons were generated (lanes 14 to 23).

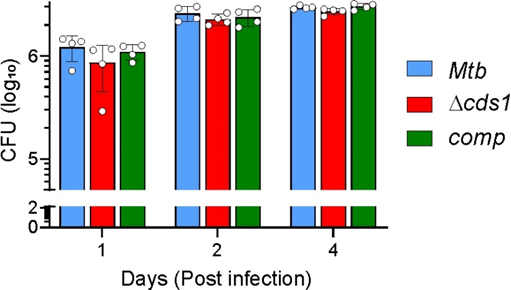

**Supplementary Figure 6. Survival of *cds1*-deficient *Mtb* in macrophages.** Peritoneal macrophages were obtained from C57BL/6 mice and infected with *Mtb* at an MOI ~0.2. Infected macrophages were lysed at indicated times after infection and lysates plated on 7H11 agar plates to determine bacillary burden. No statistically significant differences in cellular burden between WT *Mtb* and *Mtb Δcds1* cells were found. Data shown represents the mean ± SD for 4 replicates. Statistical analysis was performed using GraphPad Prism 7.02. Two-way ANOVA with Dunnett’s multiple comparisons test was used to determine statistical significance.

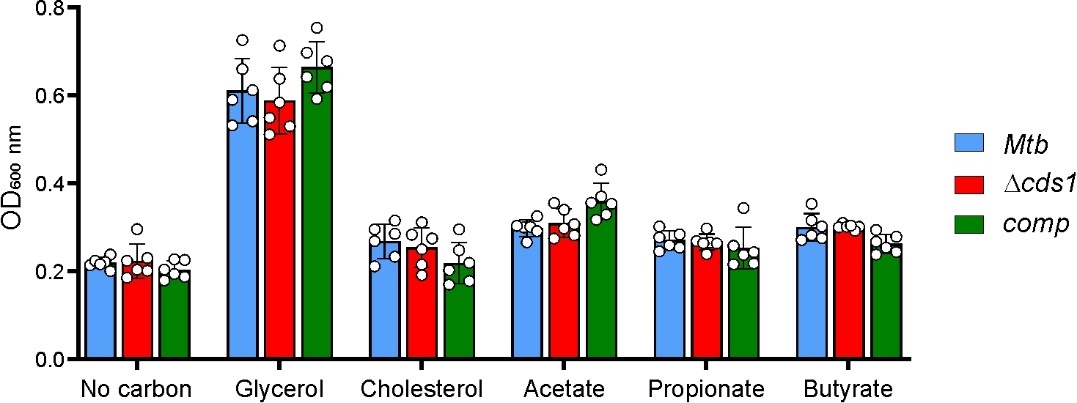

**Supplementary Figure 7. *Mtb* Δ*cds1* growth in the presence of fatty acids or precursors as a single carbon source.** Growth of *Mtb* strains was monitored at OD_600 nm_ using fatty acids or precursors (acetate) as a single carbon source. There were no significant growth differences between WT *Mtb* and Δ*cds1* cells. Data shown represents the mean ± SD for 6 replicates. Statistical analysis was performed using GraphPad Prism 7.02. Two-way ANOVA with Dunnett’s multiple comparisons test was used to determine statistical significance.

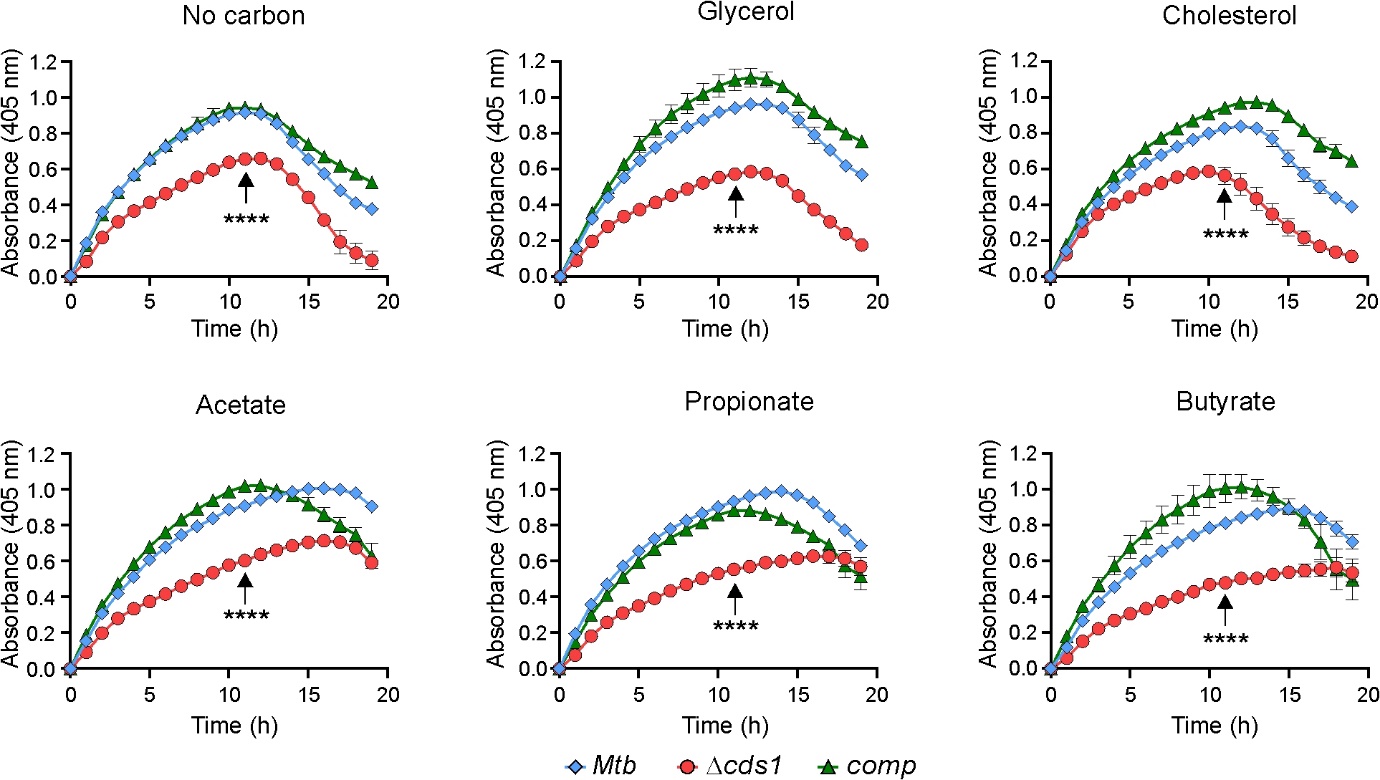

**Supplementary Figure 8. *Mtb* H_2_S production when cultured in the presence of fatty acids or precursors as a single carbon source.** The BC assay was used to measure H_2_S production of *Mtb* cultured in the presence of fatty acids or precursors (acetate) as a single carbon source. H_2_S production was significantly less in Δ*cds1* cells compared to WT or complemented cells in all growth media at 12 h (vertical arrow). Data shown represents the mean ± SEM for 4 replicates. Statistical analysis was performed using GraphPad Prism 7.02. Two-way ANOVA with Dunnett’s multiple comparisons test was used to determine statistical significance. *****P* < 0.0001.

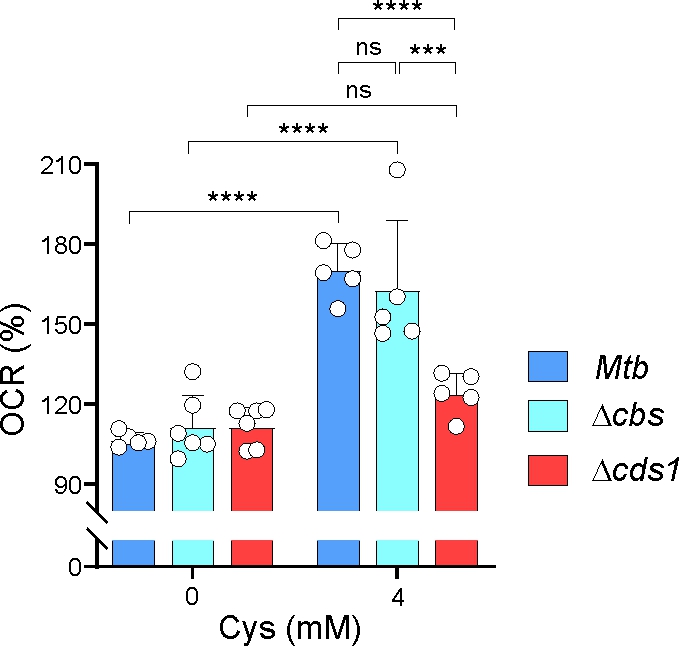

**Supplementary Figure 9. Role of Cbs (Rv1077) in *Mtb* respiration.** To determine whether Cbs (Rv1077) is important for respiration in *Mtb*, the oxygen consumption rates (OCR) of WT *Mtb*, Δ*cbs* and Δ*cds1* were measured using an Agilent Seahorse XF96 Analyzer. %OCR was measured in *Mtb* strains grown in medium containing 0 or 4 mM Cys. Cbs does not appear to play a significant role in *Mtb* respiration under these conditions. Note the role of Cds1 in respiration under the same conditions. One representative experiment is shown. Each experiment was repeated at least twice. Data represent the mean ± SD for 5 – 6 replicates. Statistical analysis was performed using GraphPad Prism 7.02. Two-way ANOVA with Tukey’s multiple comparisons test was used to determine statistical significance. ****P <* 0.001, *****P* < 0.0001, ns – statistically non-significant.

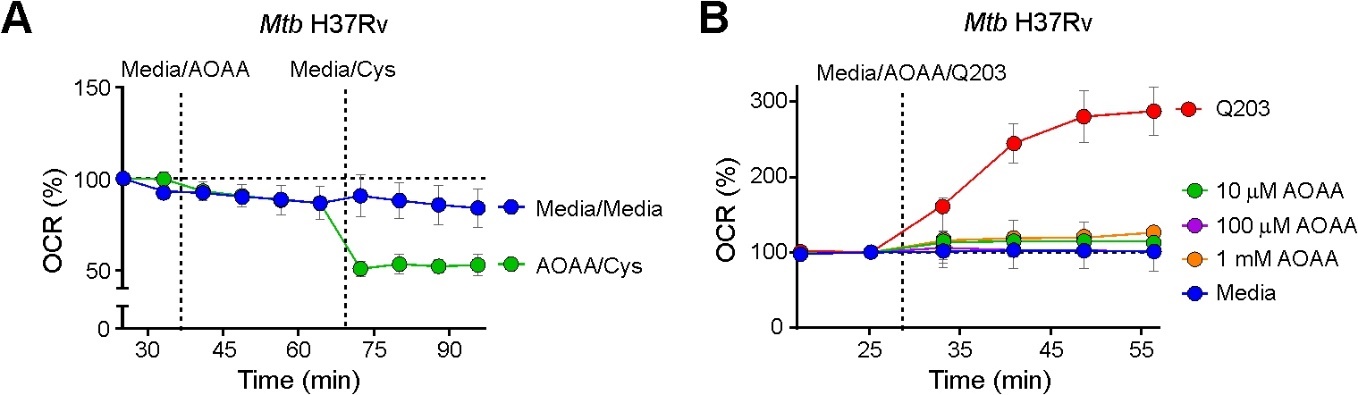

**Supplementary Figure 10. AOAA inhibits Cys-mediated increases in *Mtb* respiration.** **(A)** %OCR of *Mtb* following sequential injection of 1 mM AOAA and 1 mM Cys, or media as a control. Whereas Cys increases respiration (Fig. 5b), which can be inhibited by AOAA (Fig. 5d), the data here suggest that pre-treatment of cells with AOAA affects metabolism by targeting PLP-dependent enzymes, but not respiration. Here, following Cys addition, AOAA-treated cells are unable to respond homeostatically to Cys-generated H_2_S that normally stimulates respiration. Under these conditions, AOAA inhibits numerous PLP-dependent enzymes as well as Cds1, which collectively are unable to maintain respiratory homeostasis. **(B)** %OCR of *Mtb* grown in Cys-free medium and exposed to different concentrations of AOAA, showing that AOAA alone does not alter OCR. The 100 µM AOAA data (magenta line) is obscured by other data points. Data are representative of 2-3 independent experiments, showing mean ± SD for *n* = 6 – 8 replicates. The anti-TB drug Q203 (300x MIC_50_) that stimulates respiration was used as positive control in **(B)**.

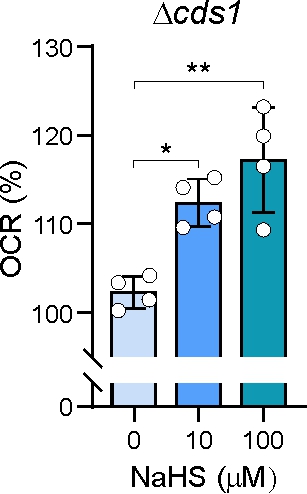

**Supplementary Figure 11. Exogenous H_2_S reverses the respiratory defect in *Mtb*** Δ***cds1* cells.** Exposing *Mtb ∆cds1* cells to different concentrations (0, 10, 100 µM) of exogenous H_2_S (NaHS) increased bacillary %OCR. Data is representative of at least two independent experiments, showing mean ± SD for *n* = 4 replicates. **P <* 0.05, ***P <* 0.01.

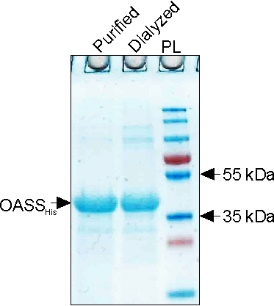

**Supplementary Figure 12.**  **SDS-PAGE of purified *O*-acetylserine sulfhydrylase (OASS).**

Recombinant OASS containing an N-terminal 6xHis tag was expressed from plasmid pET28b-EhOASS and purified from *E. coli* lysates as described previously^3^.

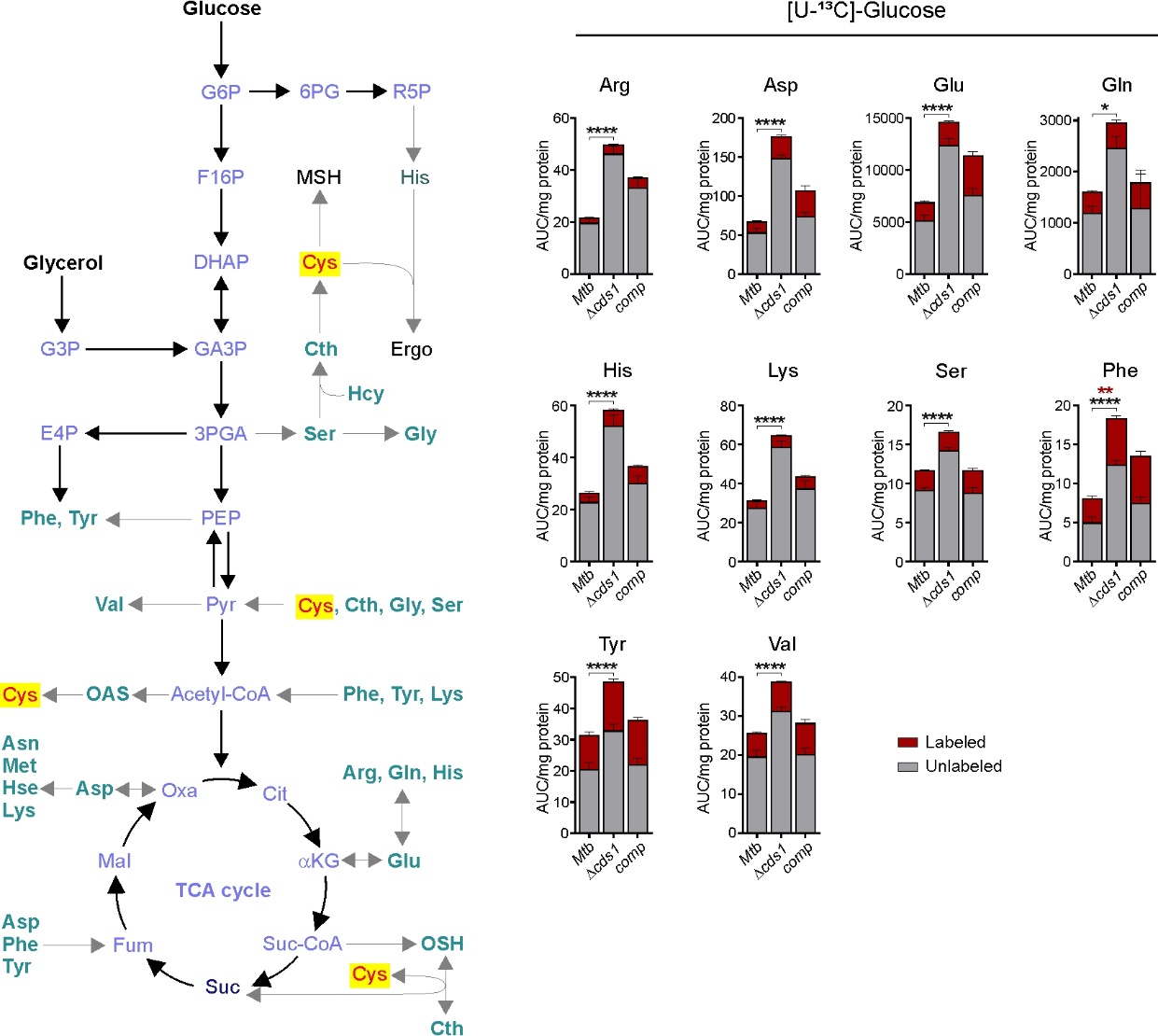

**Supplementary Figure 13. Cds1 regulates amino acid metabolism in *Mtb*.** *Mtb* strains were cultured in 7H9 medium with [U-^13^C]-Glucose (0.2%) followed by LC-MS/MS analysis. Total abundance of ^13^C-labeled (Red) and unlabeled (Gray) amino acids are indicated. Representative experiments are shown; the experiment was repeated twice. Data shown represents the mean ± SEM for 3 – 5 biological replicates. Statistical analysis was performed using GraphPad Prism 7.02. Two-way ANOVA with Dunnett’s multiple comparisons test was used to determine statistical significance. **P <* 0.05, ***P* < 0.01, *****P <* 0.0001.

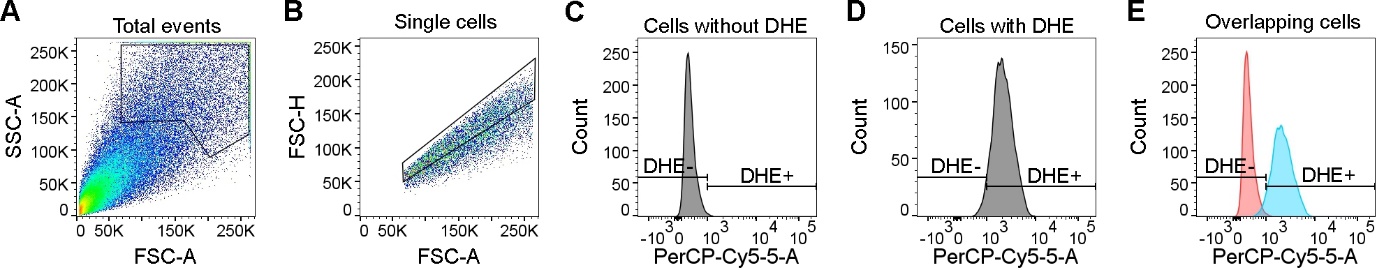

**Supplementary Figure 14. Gating strategy for detection of DHE-positive *Mtb* cells for measuring ROI.** ROI level in *Mtb* strains was measured using the dihydroethidium ROI-sensing dye (DHE, PerCP-Cy5.5). Gating strategy for **(A)** total events (100,000 cells), **(B)** single cells, **(C)** unstained cells without DHE, **(D)** DHE-stained cells, **(E)** overlapping histogram of (C) and (D) to show the population of DHE-positive and -negative cells.

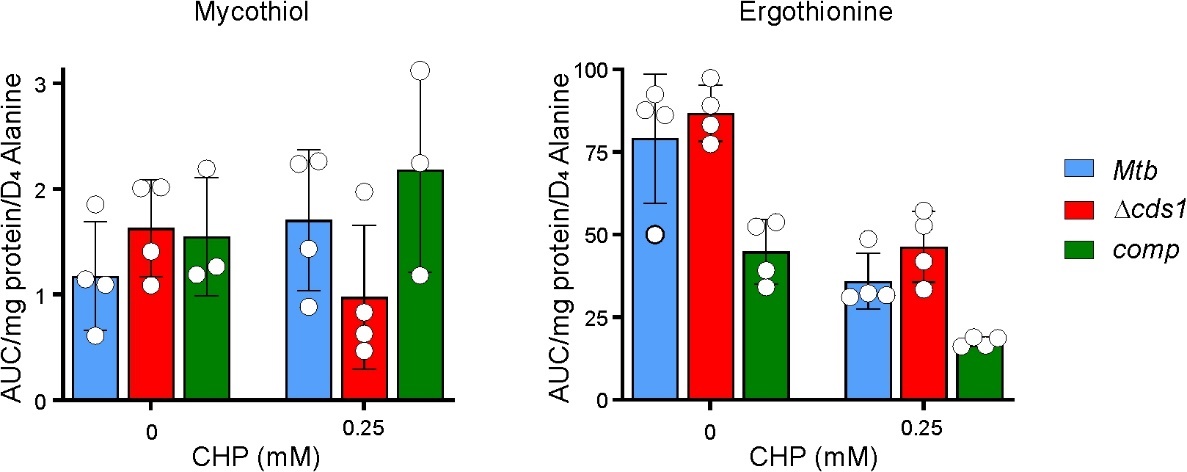

**Supplementary Figure 15. Mycothiol and ergothioneine levels in *Mtb* after exposure to CHP.** Mycothiol and ergothioneine were quantified in *Mtb* strains using LC-MS/MS after exposure to CHP for 16 h. No significant differences were observed between WT *Mtb* and *Mtb* *Δcds1* cells. Data shown represents the mean ± SEM for *n* = 3-4 replicates. Statistical analysis was performed using GraphPad Prism 7.02. Two-way ANOVA with Dunnett’s multiple comparisons test was used to determine statistical significance.

**Supplementary Table 1:** *Mtb* H37Rv enzymes putatively involved in sulfur-containing amino acid biosynthesis, H_2_S production or sulfur metabolism.

|  | **Enzymes capable of Sulfide reactions** | **Mtb Locus** | **Gene Product** | **Annotated Pathway/Function** | **Catalytic Activity** | **Ref.** |
| --- | --- | --- | --- | --- | --- | --- |
| **1** | Probable cystathionine γ-synthase/O-succinyl homoserine sulfhydrylase | Rv0391 | MetZ | Methionine Biosynthesis/ Probable Cystathionine γ-synthase | *O-*succinylhomoserine → homocysteine | ^4^ |
| **2** | Cysteine synthase | Rv0848 | CysK2 | Cysteine Biosynthesis | (1) *O-*phospho-ʟ-serine → *S-*sulfocysteine  (2) *O*-phospho-ʟ-serine + H_2_S → ʟ-cysteine + phosphate | ^5^ |
| **3** | Cystathionine β-synthase | Rv1077 | CBS | Cysteine Biosynthesis/Serine sulfhydrase/ Transulfuration Pathway | (1) homocysteine + serine → cystathionine (2) cysteine + homocysteine → cystathionine + H_2_S | ^6^ |
| **4** | Cystathionine γ-synthase/cystathionine γ-lyase | Rv1079 | MetB | Methionine Biosynthesis/Probable Cystathionine γ-synthase/ Transulfuration Pathway | (1) *O-*succinyl-ʟ-homoserine + ʟ-cysteine -> cystathionine + succinate  (2) cystathionine → α-ketobutyrate +NH_3_ | ^4^ |
| **5** | Cysteine synthase | Rv1336 | CysM | Cysteine Biosynthesis | *O*-phospho-ʟ-serine + CysO-SH → CysO-Cys + PO_4_^3-^ | ^5^ |
| **6** | Cysteine synthase | Rv2334 | CysK1 | Cysteine Biosynthesis | *O-*acetyl-ʟ-serine + H_2_S → ʟ-cysteine + acetate | ^5^ |
| **7** | Ferredoxin-dependent sulfite reductase | Rv2391 | SirA | Sulfate Assimilation | SO_3_^2-^ → S^2-^ | ^7^ |
| **8** | Methionine synthase | Rv3340 | MetC | Methionine Biosynthesis/Probable O-acetyl homoserine sulfhydrylase | (1) O-acetyl-ʟ-homoserine + methanethiol → ʟ-methionine + acetate  (2) *O-*acetyl-ʟ-homoserine + H_2_S ↔ ʟ-homocysteine + acetate | ^8^ |
| **10** | Probable cysteine desulfhydrase/  cysteine synthase /cystathionine γ-lyase? | Rv3684 | Cysteine synthase or lyase | Unclassified/Cys Metabolism/ Transulfuration Pathway? | Cysteine → pyruvate + H_2_S + NH_3_ | ^9^ |
| **11** | Probable cysteine desulfurase | Rv3025c | iscS | Carbon Sulfur Lyase | [sulfur carrier]-H + ʟ-cysteine- [sulfur carrier]-SH + ʟ-alanine | ^10, 11^ |

**Supplementary Table 2*:*** Bacterial strains used in this study.

| **Strain** | **Description** | **Source** |
| --- | --- | --- |
| *M. tuberculosis* H37Rv | Wild type (wt) | ATCC |
| *M. bovis BCG Pasteur* | Vaccine strain | ATCC |
| *M. bovis* | Wild type | ATCC |
| *M. tuberculosis* TKK-01-0027 | Clinical drug susceptible *Mtb* strain (DS27) | Alex Pym, AHRI |
| *M. tuberculosis* TKK-01-0047 | Clinical drug susceptible *Mtb* strain (DS47) | Alex Pym, AHRI |
| *M. tuberculosis* TKK-01-0035 | Clinical multi-drug resistant *Mtb* strain (MDR35) | Alex Pym, AHRI |
| *M. tuberculosis* TKK-01-0001 | Clinical multi-drug resistant *Mtb* strain (MDR01) | Alex Pym, AHRI |
| *M. tuberculosis* Δ*cbs* | *cbs* deletion mutant (*Δcbs*) in H37Rv; Hyg^R^ | This study |
| *M. tuberculosis* Δ*cds1* | *cds1* deletion mutant (Δ*cds1*) in H37Rv; Hyg^R^ | This study |
| *M. tuberculosis Δcds1::hsp_60_-cds1* | *cds1* complement (*comp*) of Δ*cds1*; Hyg^R^ and Kan^R^ | This study |
| *M. tuberculosis* CDC1551 | Wild type (wt) | ATCC |
| *M. tuberculosis Tn::rv3682* | *rv3682* transposon insertion mutant in CDC1551; Kan^R^ | John Hopkins University, School of Medicine, TARGET |
| *M. tuberculosis Tn::rv3683* | *rv3683* transposon insertion mutant in CDC1551; Kan^R^ | John Hopkins University, School of Medicine, TARGET |
| *M. tuberculosis ΔcydAB* | *cydAB* deletion mutant (*ΔcydAB*) in *Mtb* H37Rv; Hyg^R^ | Helena Boshoff^12^, NIAID |
| *M. smegmatis* mc^2^155 | Wild type (wt) | ATCC |
| *M. smegmatis wt_p_-cds1* | *rv3682-rv3683-cds1* under control of the *Mtb* native promoter (*wt_p_*) in mc^2^155 | This study |
| *M. smegmatis hsp_60_-cds1* | *rv3682-rv3683-cds1* under control of the *hsp_60_* promoter in mc^2^155 | This study |

**Supplementary Table 3:** Plasmids used in this study.

| **Vector/construct** | **Relevant genotype and properties** | **Source** |
| --- | --- | --- |
| pMV261 | *E. coli*- Mycobacterium *shuttle vector, hsp_60,_ ColE1/pAL500 oriM,* Kan^R^ | William R. Jacobs Jr. (Albert Einstein College of Medicine) |
| pET28b-OASS | Construct encoding N-terminally 6xHis-tagged EhOASS (O-acetylserine sulfhydrylase from *Entamoeba histolytica*) | Alessandro Giuffrè (CNR Institute of Molecular Biology and Pathology, Rome, Italy) |
| *cds1* phasmid | *cds1::res–hyg–res* | Michelle Larsen (Albert Einstein College of Medicine) |
| *cbs* phasmid | *cbs::res–hyg–res* | Michelle Larsen (Albert Einstein College of Medicine) |
| pMV261::*hsp_60_-cds1* | The *cds1* open reading frame cloned under the control of the *hsp60* promoter subcloned into pMV261 | This study |
| pMV261::*wt_p_-cds1* | The *rv3682-rv3683-cds1* open reading frames containing the native promoter (*wt_p_)* promoter cloned into pMV261 | This study |
| pET15b | *amp^r^, E. coli* vector used for production of his-tag fused proteins | Novagen |
| pET15b-*cds1* | *Mtb cds1* ORF subcloned into pET15b | This study |

**Supplementary Table 4:** Oligonucleotides used in this study.

| Oligonucleotide | Sequence (5’ → 3’) | Description |
| --- | --- | --- |
| Rv3684F | TATGGATCCTATGAGCGGCGGGGCCTGTATC | *cds1* forward primer for pMV261 subcloning, *Bam* HI |
| Rv3684R | GTTATCGATTAGGCTGCGGACCGCGATAC | *cds1* reverse primer for pMV261 subcloning, *Cla* I |
| ponABCF | TAAGGATCCAAGGTAGTCCGACCACGAAAC | *rv3682*, *rv3683* and *cds1* forward primer, *Bam* HI |
| ponABCR | ATAATCGATCTACCAAGCTGCGCCACAC | *rv3682*, *rv3683* and *cds1* reverse primer, *Cla* I |
| Rv3684CF | GAACCCAATGAACTATCTGAC | Forward primer for ∆*cds1* confirmation |
| Rv3684CR | GCATAGCGCATAGAGGAA | Reverse primer for ∆*cds1* confirmation |
| UUT | GATGTCTCACTGAGGTCTCT | “Universal uptag” primer for ∆*cds1* confirmation |
| Rv3684CEF | AATAATCATATGTTGAGCGGCGGGGCCT | *cds1* forward primer for pET15b subcloning, *Nde* I |
| Rv3684CER | AATAATGGATCCTCACGTCCATCGACAG | *cds1* reverse primer for pET15b subcloning, *Bam* HI |
| Rv1077CF | GGTCGACTATCGGTTGATT | Forward primer for ∆*rv1077* confirmation |
| Rv1077CR | ACATTGCGTTTATCCTCACT | Reverse primer for ∆*rv1077* confirmation |
